# Alterations to parvalbumin-expressing interneuron function and associated network oscillations in the hippocampal – medial prefrontal cortex circuit during natural sleep in App^NL-G-F^ mice

**DOI:** 10.1101/2022.02.08.479119

**Authors:** Erica S Brady, Jessica Griffiths, Lilya Andrianova, Takashi Saito, Takaomi C Saido, Andrew D Randall, Francesco Tamagnini, Jonathan Witton, Michael T Craig

## Abstract

In the early stages of Alzheimer’s disease (AD), the accumulation of the peptide amyloid-β (Aβ) damages synapses and disrupts neuronal activity and leads to disruption of neuronal oscillations associated with cognition. This is thought to be largely due to impairments in CNS synaptic inhibition, particularly via parvalbumin (PV)-expressing interneurons that essential for generating several key oscillations. Research in this field has largely been conducted in mouse models that over-express humanised, mutated forms of AD-associated genes that produce exaggerated pathology. This has prompted the development and use of knock-in mouse lines that express these genes at an endogenous level, such as the App^NL-G-F/NL-G-F^ mouse model used in the present study. These mice appear to model the early stages of Aβ-induced network impairments, yet an in-depth characterisation of these impairments in currently lacking. Therefore, using 16 month-old App^NL-G-F/NL-G-F^ mice, we analysed neuronal oscillations found in the hippocampal – medial prefrontal cortex (mPFC) during awake behaviour, rapid eye movement (REM) and non-REM (NREM) sleep to assess the extent of network dysfunction. No alterations to gamma oscillations were found to occur in the hippocampus or mPFC during either awake behaviour, REM or NREM sleep. However, during NREM sleep an increase in the amplitude of mPFC spindles and decrease in the power of hippocampal SWRs was identified. The former was associated with a decrease in the density of mPFC PV-expressing interneurons and the latter was accompanied by an increase in the synchronisation of PV-expressing interneuron activity, as measured using two-photon Ca^2+^ imaging. Furthermore, although changes were detected in local network function of mPFC and hippocampus, long-range communication between these regions appeared intact. Altogether, our results suggest that these NREM sleep-specific impairments represent the early stages of circuit breakdown in response to amyloidopathy.

## Introduction

Alzheimer’s disease (AD) is a progressive neurodegenerative disorder and the most prevalent form of dementia worldwide^1^. In the preclinical stages of the disease, preceding overt signs of cognitive decline, the build-up and aggregation of the peptide amyloid-β (Aβ) causes detriment to the synapse and neuronal activity, before accumulating further to form the extracellular plaque pathology that characterises AD. In turn, these detriments disrupt the synchronous communication between groups of neurons that generates the neuronal oscillations associated with cognitive function^2–7^. These impairments to the electrical activity of the brain can be measured in humans as hyperactivity of the hippocampus and areas of the default mode network (DMN)^8,9^ and lead to the generation of epileptiform activity^10^ and a general “slowing” of the EEG^11^. Evidence suggests that altered inhibitory neurotransmission underlies this breakdown in circuit function^12^. In particular, parvalbumin (PV)-expressing interneurons have been shown to exhibit both hypo- and hyperactivity in mouse models of amyloidopathy^13–15^, with the former being linked to a reduction in the amplitude of gamma oscillations and the emergence of epileptic spikes.

Sleep disturbances, particularly disruptions to the quality and duration of non-rapid eye movement (NREM) sleep, are also a common feature during the early stages of AD, and are strongly linked to cognitive decline^16,17^. Importantly, the long-term consolidation of memory is thought to occur during NREM sleep, with significant research emphasis being placed on the activity of the hippocampal – medial prefrontal cortex (mPFC) circuit, given its prominent roles in episodic and spatial memory, as well as higher-order executive functions^18,19^. The generation and temporal coupling of three cardinal neuronal oscillations – the cortical slow wave oscillation (SWO), spindles, and hippocampal sharp-wave ripples (SWR) – facilitates the reorganisation of newly encoded hippocampal memories into distributed cortical networks for long-term storage, in a process called systems consolidation^20,21^. Perturbation of the SWO and spindles is observed in humans with AD^22,23^, with similar disruptions to the SWO occurring in mouse models of amyloidopathy^24–26^, alongside impairments to SWRs^27^.

Understanding neuronal network dysfunction in the preclinical stages of AD is imperative for the identification of early-detection biomarkers and for the development of treatments to be delivered prior to the occurrence of irreversible neurodegeneration and cognitive decline. However, the majority of research to date has been conducted in transgenic mouse models that develop exaggerated pathology, comparative to the later stages of the disease. This is due to the over-expression of humanised, mutated AD-linked genes that are co-expressed alongside murine homologs^28^. Indeed, the widespread use of such over-expression models may provide one explanation why experimental pharmaceutical interventions that show efficacy in mice have commonly failed in phase III clinical trials^28^. This has prompted the innovation of knock-in lines that have had the endogenous mouse gene replaced with the human homolog, thus achieving physiological levels of expression^28–30^. Conducting experiments in knock-in mouse models that are not confounded by non-physiological over-expression of mutated genes should provide researchers with a greater understanding of how early Aβ pathology alters brain physiology. With this in mind, we analysed the neuronal oscillatory activity of the hippocampal-mPFC circuit in sixteen-month-old App^NL-G-F/NL-G-F^ mice across three different brain states: awake behaviour, rapid-eye movement (REM) and NREM sleep. Our goal was to assess Aβ-induced network dysfunction before cognitive decline, and to determine whether this dysfunction displayed brain region- and state-specific effects.

## Methods

### Animals

Male and female, App^NL-G-F/NL-G-F^ mice (APP) at sixteen-months-old (originally obtained from the MRC Harwell Institute, bred internally, and maintained on a C57BL/6J background) and wildtype (WT) littermate controls were used in this study^30,31^. All animals were housed on a 12-hour light-dark cycle at a temperature of 22 ± 2 °C and humidity of 45 ± 15% and had *ad libitum* access to food and water. For electrophysiological studies, N = 6 mice per genotype were implanted with electrodes but due to post-operative complications, N = 5 mice per genotype were used for experiments. N = 7 WT and N = 8 APP were used for immunohistochemical analysis. For two-photon imaging, APP mice were crossbred with mice expressing Cre recombinase in parvalbumin-expressing interneurons (PV-cre mice; The Jackson Laboratory, #008069) or somatostatin-expressing interneurons (SST-cre mice; The Jackson Laboratory, #013044) to generate App^HOM^::PV-cre (PVAPP) and App^WT^::PV-cre (PVWT) mice, or App^HOM^::SST-cre (SSTAPP) and App^WT^::SOM-cre (SSTWT), respectively. N = 3 PVWT, N = 5 PVAPP, N = 2 SSTWT and N= 4 SSTAPP mice were used for experiments. All procedures were carried out in accordance with the UK Animals (Scientific Procedures) Act 1986 and EU Directive 2010/63/EU and were reviewed by the University of Exeter Animal Welfare and Ethical Review Body.

### Surgical implantation of electrodes

Surgical procedures were performed using aseptic techniques. APP and WT mice were anesthetised with isoflurane and placed in a stereotaxic frame (Stoelting, IL, USA). An incision was made in the scalp to expose the skull and microscrews (Antrin, CA, USA) were inserted into the frontal, parietal and interparietal plates bilaterally to anchor the implant to the skull. 1 mm diameter craniotomies were made over mPFC (in mm, from Bregma: AP = +1.75, ML = +0.25) and dorsal CA1 (in mm, from Bregma: AP = −2.0, ML = +1.4) and four-channel electrodes (Q1×4-5mm-200-177-CQ4, Neuronexus, MI, USA) were implanted to a depth (in mm, from the brain surface) of 1.7 (mPFC) and 1.5 (CA1), respectively. Silver wire (World Precision Instruments, FL, USA) and silver conductive paint (RS Components, Corby, UK) were used to connect the ground channel of each electrode array to an anchor screw overlying the cerebellum. Light-curable dental cement (RelyX Unicem, Henry Schein, NY, USA) was used to secure the electrodes to the skull and anchor screws, and the scalp was closed around the base of the implant using a suture. Subcutaneous carprofen (5 mg/kg) was used for the management of post-operative pain as required.

### Electrophysiology data acquisition

Neural signals were acquired by connecting each electrode array to an Omnetics adapter (OM26, NeuroNexus) and headstage amplifier (RHD2000, Intan Technologies, CA, USA). The two headstages were connected to a dual headstage adapter (Intan) that allowed signals to be relayed along a single tether cable (Intan RHD SPI) to an OpenEphys acquisition board. Signals were bandpass filtered online at 0.1-7932 Hz and continuously recorded at 30 kHz. Signals recorded in CA1 in two mice (1 APP; 1 WT) were of poor quality (potentially due to poor grounding) and so were excluded from analysis.

Recordings commenced 8 weeks after surgery. Animals were recorded in their home cage for three hours during the lights-on circadian epoch to enable recording of neural signals during sleep. An overhead light remained on throughout recording (440 lux). A Logitech colour webcam was fitted overhead (30 Hz frame rate) and videos were recorded throughout neural signal acquisition. To record neural activity during ambulation, animals were placed in a square open-field and allowed to explore for 1 hour. Two light-emitting diodes (LEDs) were attached and grounded to one of the headstages that allowed the animal’s position to be tracked using the software Bonsai^32^. Tracking data collected from Bonsai were acquired through the OpenEphys recording system at the same time as the neural signals. To allow for the LEDs to be visualised, all lights were switched off apart from an overhead LED light (1 lux).

### Verification of electrode locations

At the end of experimental procedures, mice were deeply anaesthetised using an overdose of sodium pentobarbital (300 mg/kg) and electrolytic lesions were made at electrode sites. Mice were then transcardially perfused with 0.1 M phosphate buffer (PB) followed by 4% paraformaldehyde (PFA) in 0.1 M PB. Brains were extracted, postfixed in 4% PFA in 0.1 M PB overnight, and cryoprotected in 30% w/w sucrose in 0.01 M phosphate buffered saline (PBS). Brains were cut into 40 μm coronal sections using a freezing microtome (SM2010R, Leica Microsystems) and every fourth slice through mPFC and dorsal CA1 was mounted onto a slide and stained with DAPI. Sections were visualised using an epifluorescence microscope (Nikon Eclipse E800) and recording locations were confirmed from the lesions and electrode tracts. mPFC regions were defined based on previous literature^33^, with electrode positions found in both the anterior cingulate cortex (ACC) and prelimbic (PL) cortex (***Supplementary Figure 1A, B***). Hippocampal electrode placement was identified through histology and the changing polarity of SWRs in local field potential (LFP) traces recorded through the layers of CA1^34^. Only electrodes located in *Stratum pyramidale* (*Str.P*) were used for analysis (***Supplementary Figure 1C, D***).

### Viral injection surgery and two-photon imaging

PVWT, PVAPP, SSTWT and SSTAPP mice underwent similar surgical procedures as described above. The skull was exposed, and a craniotomy was made over dorsal CA1 (in mm, from Bregma: AP = −2.2, ML, +2.0). 250 nl of AAV9/2-CAG-dlox-GCaMP6f-dlox-WPRE-SV40 (5×10^12^ GC/ml) (VVF, University of Zurich) was infused into the CA1 region at a depth of 1.3 mm from the dura (rate: 50 nl/min) using a pulled glass micropipette and microinjection pump (World Precision Instruments). Three weeks after viral injection, mice were implanted with an optical cannula overlying dorsal CA1, as previously reported^35^. Briefly, anaesthesia was induced using ketamine (73 mg/kg)/medetomidine (0.44 mg/kg) and maintained using isoflurane. The skull was exposed, and a 3 mm diameter craniotomy was drilled, centred on the previous injection site. The dura was removed, and a column of cortex was aspirated whilst continuously irrigating with chilled cortex buffer (in mM: NaCl 125, KCl 5, glucose 10, HEPES 10, CaCl_2_ 2, MgSO_4_ 2). A stainless-steel cannula (3 mm o.d.; 2.4 mm i.d.; 1.5 mm long) with a 3 mm diameter coverslip (Warner Instruments) affixed to one end (using NOA71 adhesive, Norland) was lowered into the craniotomy to the depth of the external capsule and was glued in position (Loctite). The skull was then sealed with dental cement (Simplex Rapid, Kemdent) and a metal head-fixation bar was affixed to the implant.

Recordings were performed under light (0.5-1%) isoflurane anaesthesia 150-240 min after initial induction of anaesthesia for surgery. Mice were transferred to a two-photon microscope and the head-fixation bar was secured using a clamp. Body temperature was maintained at 37°C using a homeothermic blanket. A piezoelectric transducer beneath the thorax continuously monitored breathing rate as a proxy measure of anaesthetic depth, and the isoflurane dose was adjusted to maintain a consistent breathing rate between 150-200 breaths per minute. Images were acquired at 16x objective magnification (Nikon Plan Fluorite, 0.8 NA, 3 mm WD) using a microscope (Hyperscope, Scientifica) equipped with a Ti:sapphire pulsed laser (Chameleon Discovery, Coherent) and galvanometric scan mirrors. GCaMP6f was excited at 920 nm (power <50 mW at the sample) and fluorescence was collected using a 500-550 nm emission filter. Images were acquired at 30 Hz using ScanImage 4.0 software^36^. Spontaneous GCaMP6f signals were recorded continuously for 300 s. Signals were captured 200-300 μm beneath the external capsule in PVWT and PVAPP mice (around the depth of CA1 *Str.P*) and 100-200 μm beneath the external capsule in SSTWT and SSTAPP mice (around the depth of CA1 *Str. Oriens*). Several regions of interest (ROIs) (4.5 (3 7) [Med (Min Max)]) containing multiple GCaMP6f-positive neurons (5 (2 16) [Med (Min Max)]) were imaged per mouse.

### Electrophysiology data analysis

All analysis was performed using custom-made scripts in MATLAB (Mathworks).

#### Detection of sleep states and sleep-associated neuronal oscillations

Periods of NREM^37,38^ and REM sleep were isolated from signals using an algorithm based on previously published work^39,40^. LFP signals were first split into 30 second epochs and a theta:SWO/delta (6-12 Hz:1-4 Hz) power ratio was created. If the power ratio in a 30 second epoch fell below a threshold (median + 1 SD), and was also accompanied by a lack of movement, the epoch was considered sleep. If an epoch did not meet this requirement, it was considered to be awake. These signals were put through the same algorithm using 10 second bins to further split sleep epochs into NREM and REM sleep (***Supplementary Figure 2A***).

To analyse the dynamics of each of the three cardinal oscillations, custom-made detection scripts based on previously described methods were used to isolate and analyse individual oscillation cycles^37,41–48^. The SWO was detected by first low-pass-filtering the raw SWS signal (1.5 Hz) and all negative to positive zero-crossings were found. Two consecutive negative-positive zero-crossings within 0.25-2 s (0.5-1.25 Hz) of each other and the largest peak and trough between them were used in the next stage of detection. Events were classed as a SWO if the peak was greater than the 60th percentile of all detected peaks and the total amplitude (peak-trough) was also greater than the 60th percentile of all detected amplitudes.

For the detection of spindle (9-16 Hz) and ripple (130-250 Hz) events, the raw SWS signal was first bandpass filtered, z-scored and peaks were interpolated using cubic spline interpolation. Peaks that crossed a 3.5 SD threshold were identified and start and stop times of 1.5 SD either side were taken for the detection of spindles. A 6.5 SD threshold with 2 SD start and stop times were used for the detection of ripples. Events were then classed as spindles if they were between 0.5 and 2 s long and had an intra-spindle frequency of 11-15 Hz and as ripples if their duration was between 30-250 ms and had an intra-ripple frequency of 130-250 Hz.

#### Oscillation coupling

Analysis windows spanning 2 s on each side of detected Down states were used to analyse the phase-amplitude coupling (PAC)^49^ between detected SWO events and gamma oscillations. Spindles were considered coupled to the SWO If the peak of a detected spindle event occurred within 1 s after the peak of a detected Down state. Sections of raw signal that began 2 s before the onset of the Down state and ended 2 s after the peak of the spindle event were used to analyse their PAC. Similar analysis methods were applied for the SWO and ripples, except a 0.5 s period after the ripple peak was used for PAC. If a ripple began within 2 s following the onset of a spindle, they were considered coupled.

#### Open Field Power Analysis

The two LEDs attached to the headstage were used to track the animal’s position. Time-points in which the LEDs failed to reach detection levels or occurred out of range of the OF were automatically rejected and adjoining data points linearly interpolated. Speed was initially computed in 1 s time-bins ranging from 1-30 cm/s using the position of the animal. The LFP was then split into corresponding 1 s time bins and the total power (area under the curve, trapezoid method) was found for theta (6-12 Hz), low gamma (30-60 Hz) and high gamma (60-120 Hz) oscillations. Spectral analysis was computed using the Chronux toolbox^50^, using the function *mtspecgramc*. 1 s time bins with 50% overlap were used as well as 3 tapers, with the resultant spectra normalised to total power. Gamma oscillations were defined based on identified peaks in the spectra and electrical noise at 50 Hz and 100 Hz (4^th^ order IIR Butterworth Bandstop, 47-53 Hz and 97-103 Hz, respectively) was removed from the spectra for analysis.

### Two-photon imaging analysis

Custom routines in MATLAB and FIJI software ^51^ were used. Mechanical drift was corrected by cross-correlation-based subpixel registration^52^ to the average of the first second (30 frames) of each image time series. Sum projection images were then used to manually draw ROIs around visually identified cell bodies, and cellular fluorescence time series, F_ROI_(t), were constructed by averaging the ROI pixels in each frame. A correction for neuropil contamination was applied, following F_ROI_(t) = F_ROI__measured(t) – F_ROI__neuropil(t) x R^53^. The neuropil signal for each ROI (F_ROI__neuropil) was calculated by averaging the pixels within a 2 μm annulus expanded around each ROI, excluding pixels within other cellular ROIs. R was the linear regression coefficient between the measured ROI signal and its neuropil signal. Slow drifts in the fluorescence time series were corrected by subtracting the 8th percentile in a 30 second window centred on each time point, and relative fluorescence time series (ΔF/F) were calculated as (F_ROI_(t)-F0)/F0, where F0 was the median of F_ROI_(t) after drift correction. To detect fluorescence transients, we calculated the SD of the 20-80th percentile of ΔF/F, and identified contiguous samples that exceeded F0 + 2SD and had a peak amplitude of at least F0 + 7SD. For correlation analysis, we calculated, for each cell, the mean pairwise zero-lag cross correlation between the cell’s ΔF/F trace and the ΔF/F traces of all other simultaneously recorded cells.

### Immunohistochemistry

Animals were transcardially perfused and brains were removed and sectioned, as described above. Sections were first washed in PBS with 0.3% Triton X-100 (PBS-T) before being immersed in 100 mM glycine in PBS-T. Sections were then washed in PBS-T and incubated in blocking solution (3% goat serum in PBS-T) for 1 hour. Primary antibodies (rabbit anti-PV, 1:5000, Swant or rabbit anti-Aβ_1-42_, 1:1000, Cell Signalling Technology) were diluted in blocking solution, applied to the sections, and incubated overnight at 4°C. The next day, sections were washed in PBS-T and incubated in blocking solution containing biotinylated goat anti-rabbit secondary antibody (1:250, Vector Laboratories) for 1 hour at room temperature. Sections were then rinsed in PBS-T and incubated in blocking solution containing DyLight 488 Streptavidin (1:250, Vector Laboratories) for 1 hour. Finally, sections were washed, mounted onto glass slides, and covered with glass cover slips with mounting medium containing DAPI.

Images were acquired at 10x magnification on a Leica confocal laser scanning microscope. Only the right hemisphere was imaged and each brain region was sampled at a similar distance from bregma between animals. Brain regions were defined according to published criteria^33^ and the Allen Brain Atlas (https://mouse.brainmap.org/). To quantify the number of PV-expressing interneurons, the cell counter plugin was employed using FIJI software^51^. All cell counts were normalised to the brain region area. Widespread Aβ_1-42_ pathology was observed in both the mPFC and hippocampus in APP mice (***Supplementary Figure 3***).

### Statistical analysis

For two-photon imaging analysis, cell was used as the experimental unit (N) as amyloidopathy is a progressive pathology that affects different cells at different times, often dependent on proximity to Aβ aggregates^3^. Nonetheless, cells within individual animals are not fully independent and may covary^54^. To control for the nesting of cell within animal, the data were modelled using generalised linear mixed effects models (GLMMs) that included ‘animal’ as a random factor, allowing by-animal random intercept and by-animal random slope (R, lme4 package). Event frequencies were expressed as the total number of events in 300 s and modelled in a GLMM with Poisson distribution and log link function. Event amplitudes were modelled in a GLMM with lognormal distribution to account for positive skew in the population distributions. Cellular correlations fit a normal distribution and were modelled in an LMM. The effect of genotype was assessed via likelihood ratio chi-squared test.

All other statistical testing was carried out using Prism 9 (GraphPad). Normality of the data was tested using a D’Agostino and Pearson test. A Mann-Whitney U-test of comparable ranks was used for nonparametric data. All non-parametric results are displayed as box plots showing the median, range and the inter-quartile range (IQR).

### Data availability

All raw data that supports the findings herein are available from the corresponding author upon reasonable request.

## Results

As disruptions to the sleep cycle appear in both humans with AD and mouse models of amyloidopathy^55^, it is important to first state that no statistically significant difference was found between genotypes in the time spent in awake states, NREM or REM sleep (***Supplementary Figure 2B***).

### The power of gamma oscillations is not altered across brain states

Gamma oscillations (30-120 Hz) group the firing of neurons into distinct epochs to enable precise encoding and retrieval of information^56,57^. They can be found nested within theta oscillations during awake behaviour and REM sleep and within the cortical SWO during NREM sleep^58,59^. Impairments to gamma oscillations have been identified during wake in humans with AD^60,61^ and in several mouse models of amyloidopathy^13,14,24,25,62,63^ along with the dysfunction of the inhibitory interneurons^13–15^ that drive them. We therefore hypothesised that similar impairments would be present in the mPFC and CA1 region of the hippocampus of APP mice during awake behaviour and in different sleep stages. However, in the mPFC no statistically significant change to the power of low (30-60 Hz) or high (60-120 Hz) gamma oscillations was identified in APP mice relative to WT when mice were ambulating at slow speeds (< 5 cm/s) or high speeds (> 20 cm/s) (***Figure 1A-C***). There was also no difference between genotypes in low and high gamma oscillation power during REM sleep (***Figure 1B-E***) and during SWO-nested gamma oscillations in NREM sleep (***Figure 1F-H***).

**Figure 1.**
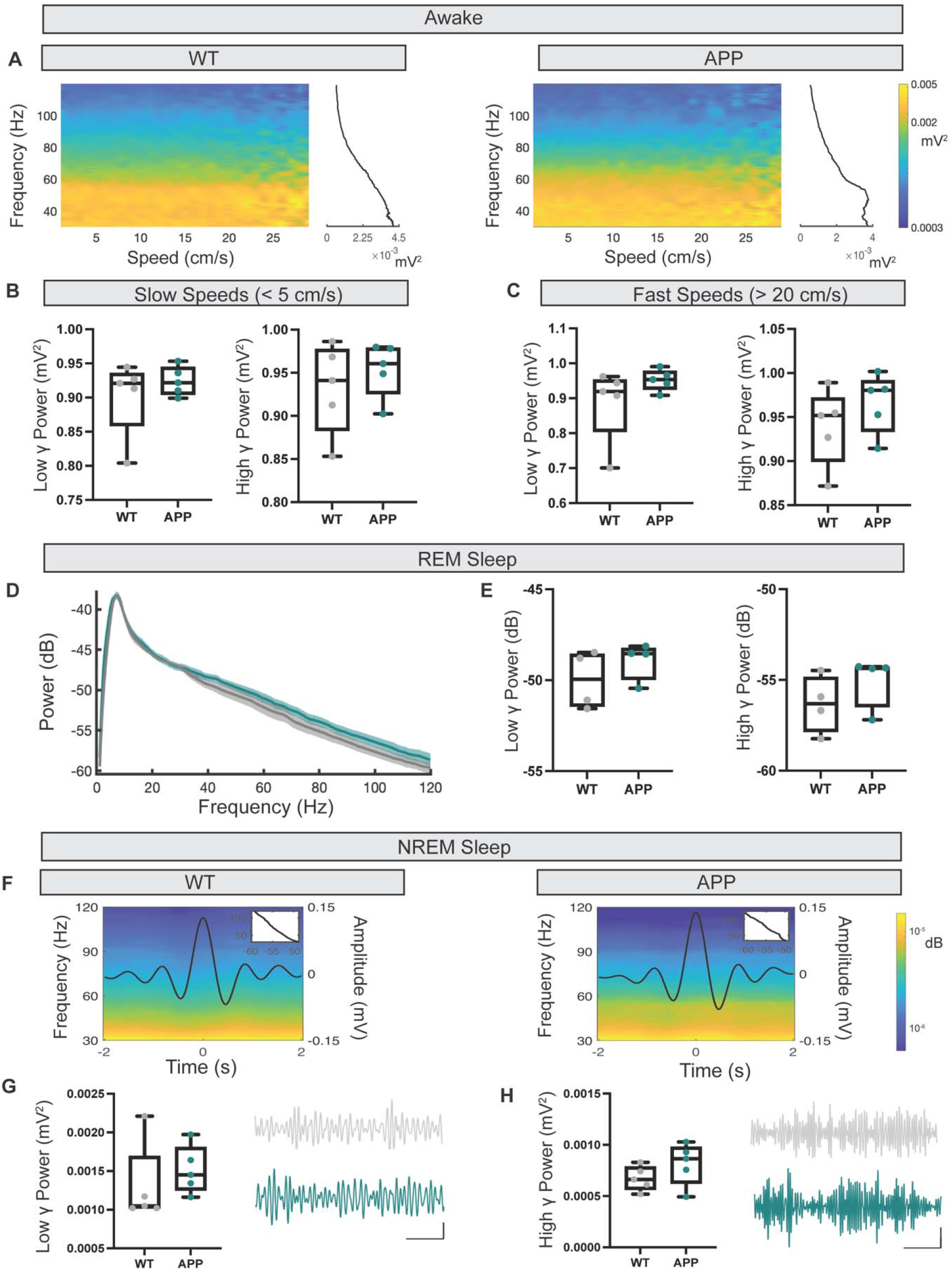
No deficits to gamma oscillatory power in mPFC. **A** Example spectrograms displaying a speed-modulated low and high gamma power in WT and APP mice. **B** No statistically significant difference was found to the power of low gamma (WT: 0.920 (0.859 – 0.936) vs APP: 0.922 (0.905 – 0.945) mV^2^, U = 11, p = 0.84, Mann-Whitney U test) and high gamma (WT: 0.941 (0.883 – 0.977) vs APP: 0.961 (0.926 – 0.979) mV^2^, U = 10, p = 0.69, Mann-Whitney U test) at slow ambulatory speeds. **C** No statistically significant difference was found to the power of low gamma (WT: 0.920 (0.804 – 0.953) vs APP: 0.954 (0.925 – 0.978) mV^2^, U = 6, p = 0.22, Mann-Whitney U test) and high gamma (WT: 0.952 (0.899 – 0.972) vs APP: 0.980 (0.936 – 0.992) mV^2^, U = 8, p = 0.42, Mann-Whitney U test) at fast ambulatory speeds. **D** Average power spectra ± SEM of both WT (grey) and APP (turquoise) CA1 LFPs during REM sleep. **E** No statistically significant difference was found to the power of low gamma (WT: – 49.94 (−51.45 – −48.55) vs APP: −48.54 (−49.97 – −48.23) dB, U = 4, p = 0.34, Mann-Whitney U test) and high gamma (WT: −56.31 (−57.85 – −54.84) vs APP: −54.36 (−56.49 – −54.29) dB, U = 3, p = 0.20, Mann-Whitney U test) during REM sleep. **F** Example average SWO traces (black lines) overlaid on the corresponding LFP power spectrum showing the power of SWO-locked gamma oscillations for both WT and APP mice. Indented plots (top right corner) display the mean gamma power over the time-course of the LFP power spectrum. **G-H** No statistically significant difference was found to the power of low (WT: 0.0010 (0.0010 – 0.0017) vs APP: 0.0014 (0.0013 – 0.0018) mV^2^, U = 6, p = 0.2, Mann-Whitney U test) and high gamma (WT: 0.00066 (0.00057 – 0.00079) vs APP: 0.00087 (0.00063 – 0.00098) mV^2^, U = 6, p = 0.22, Mann-Whitney U test). Box plots show median, IQR and ranges. Descriptive statistics display median and IQR.

Consistent with these results, no statistically significant change in gamma oscillatory power was found in CA1 during awake behaviour at slow and fast ambulatory speeds (***Figure 2A-C***) or in REM sleep when comparing APP with WT mice (***Figure 1D-E***). Analysis of hippocampal SWO-nested gamma oscillations within NREM sleep was omitted from this analysis, as the function of these oscillations in this region are still to be elucidated. Despite no statistically significant power differences, a trending decrease in the power of low and high gamma oscillations can be seen in APP mice compared with WT across brain states, potentially indicating the beginning of gamma oscillatory deficits at this pathological stage. These results raised the question as to whether the dysfunction of other neuronal oscillations precedes gamma breakdown, which prompted us to focus our attention on the neuronal oscillations underlying memory consolidation in the hippocampal-mPFC circuit during NREM sleep.

**Figure 2.**
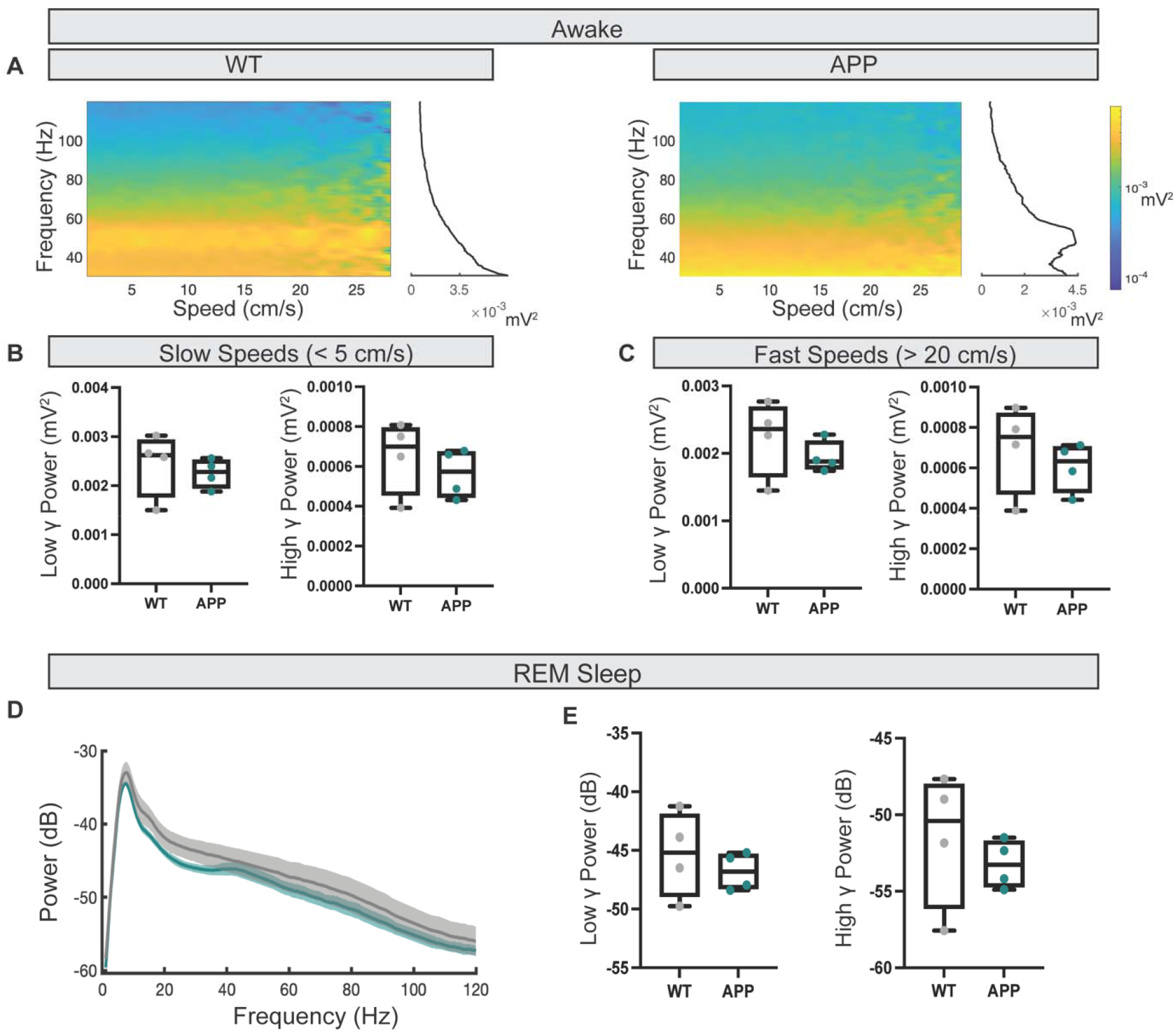
No deficits to gamma oscillatory power in CA1. **A** Example spectrograms displaying speed-modulated low and high gamma oscillation power in WT and APP mice. **B** No statistically significant difference was found to the power of low gamma (WT: 0.0020 (0.0017 – 0.0029) vs APP: 0.0023 (0.0020 – 0.0025) mV^2^, U = 4, p = 0.34, Mann-Whitney U test) and high gamma (WT: 0.0006 (0.0004 – 0.0007) vs APP: 0.0006 (0.0004 – 0.0007) mV^2^, U = 6, p = 0.68, Mann-Whitney U test) at slow ambulatory speeds. **C** No statistically significant difference was found to the power of low gamma (WT: 0.0024 (0.0017 – 0.0027) vs APP: 0.0018 (0.0017 – 0.0022) mV^2^, U = 5, p = 0.49, Mann-Whitney U test) and high gamma (WT: 0.0007 (0.0005 – 0.0009) vs APP: 0.0006 (0.0005 – 0.0007) mV^2^, U = 4, p = 0.34, Mann-Whitney U test) at fast ambulatory speeds. **D** Average power spectra ± SEM of both WT (grey) and APP (turquoise) CA1 LFPs during REM sleep. **E** No statistically significant difference was found to the power of low gamma (WT: −45.19 (−48.95 – −41.90) vs APP: −46.81 (−48.28 – −45.31) dB, U = 6, p = 0.67, Mann-Whitney U test) and high gamma (WT: −50.41 (−56.14 – −48.00) vs APP: −53.27 (−54.71 – −51.71) dB, U = 5, p = 0.49, Mann-Whitney U test) during REM sleep. Box plots show median, IQR and ranges. Descriptive statistics display median and IQR.

### Decreased ripple power and increased coordination of PV-expressing interneuron activity in the hippocampus in APP mice

Consolidation of newly acquired hippocampal memories occurs during NREM sleep as a result of the temporal communication between the cortical SWO, spindles and hippocampal SWR^20,21^. SWR are oscillatory events that coincide with the replay of previously encoded information^64,65,66^ (***Figure 3A***), with disruptions to these oscillations impairing the consolidation of hippocampus-dependent memories^67^. As SWR are impaired in over-expression mouse models of amyloidopathy^27^, we hypothesised that disruptions to SWR would also be seen in APP mice. A statistically significant decrease in the power of detected ripple events was found in APP mice (***Figure 3B***), yet interestingly, this was not accompanied by a change in SWR amplitude (***Figure 3Cii***). Additionally, a trend towards an increase in the occurrence of detected ripple events was found (***Figure 3Civ***), yet there was no change in the duration and frequency of detected ripple events in APP animals compared with WT (***Figure 3Ci, iii***).

**Figure 3.**
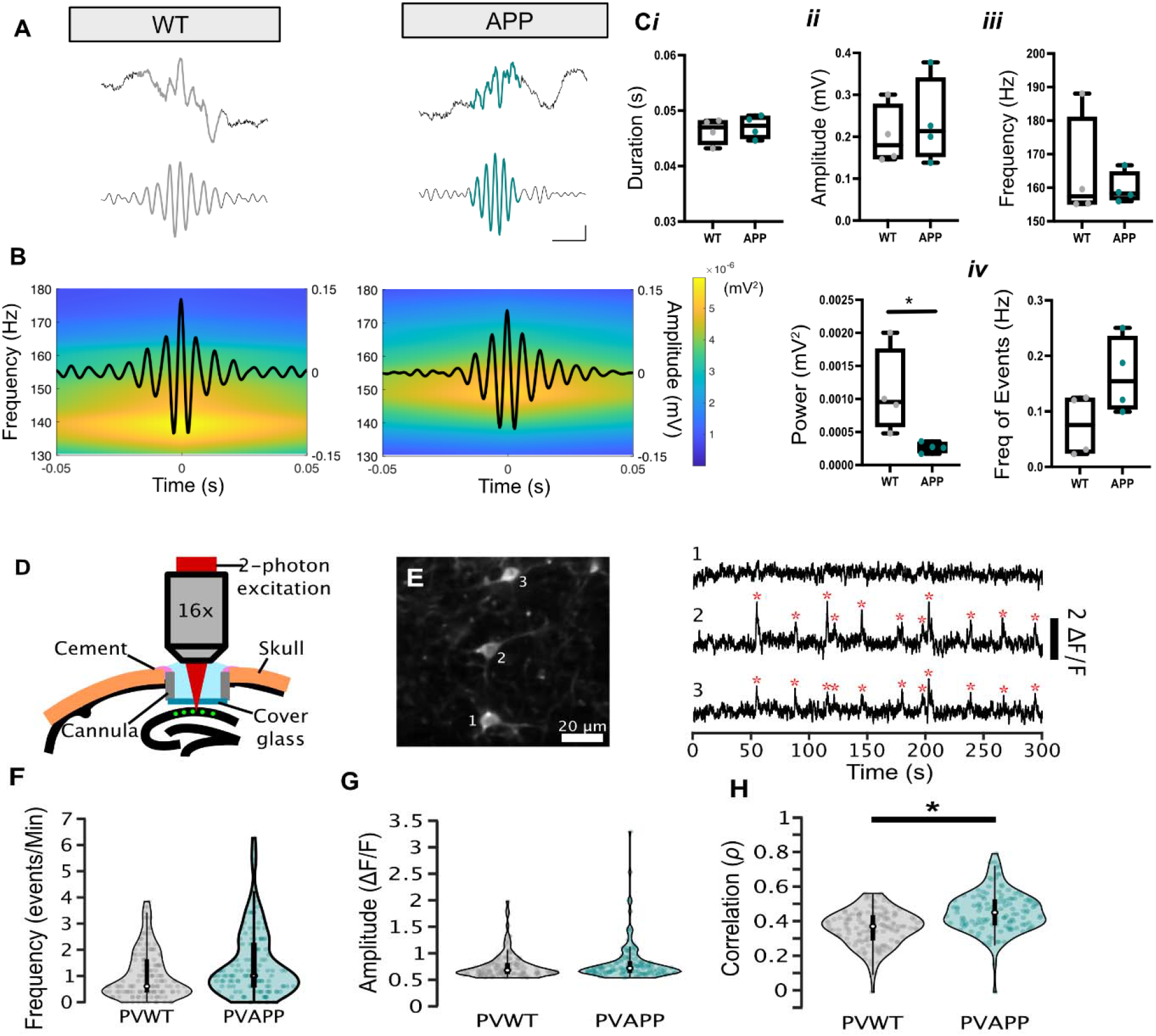
Decreased power of hippocampal ripple events and increased coordination of PV-expressing interneuron activity in APP mice. **A** Example raw (top) and 130 – 250 Hz bandpass filtered (bottom) traces of detected ripple events from WT and APP mice (coloured segments). Scale bars: 20 ms, 50 μV (bottom), 100 μV (top). **B** Example average ripple traces (black lines) overlaid on corresponding LFP power spectra showing average ripple power. A statistically significant decrease in the average ripple power can be seen in APP mice compared with WT controls (WT: 0.00095 (0.00059 – 0.00175) vs APP: 0.00027 (0.00019 – 0.00034) mV^2^, U = 0, p = 0.02, Mann-Whitney U test). **C** No statistically significant difference was found in the average ripple duration (WT: 0.050 (0.044 – 0.048) vs APP: 0.047 (0.045 – 0.049) s, U = 5, p = 0.48, Mann-Whitney U test) ***(i)***, amplitude (WT: 0.18 (0.15 – 0.28) vs APP: 0.21 (0.15 – 0.34) mV, U = 7, p = 0.88, Mann-Whitney U test) ***(ii)*** and frequency (WT: 157.4 (155.2 – 180.9) vs APP: 158.3 (156.5 – 164.7) %, U = 7, p = 0.88, Mann-Whitney U test) ***(iii)***. A trending increase in the occurrence of detected ripple events was found in APP mice compared with WT controls (WT: 0.08 (0.03 – 0.12) vs APP: 0.15 (0.10 – 0.23) Hz, U = 3, p = 0.2, Mann-Whitney U test) ***(iv).*** **D** Schematic of two-photon imaging of PV+ neurons (green) in CA1 of the dorsal hippocampus. **E** Left: Two-photon image of three representative GCaMP6s-expressing cells in a PVWT mouse. Right: GCaMP6s fluorescence traces (ΔF/F) recorded from these cells. Red stars denote detected spontaneous Ca^2+^ events. **F-G** No statistically significant change was found in PVAPP cells (N = 128 cells, 28 (13 41) cells/mouse [Med (Min Max)]) compared with PVWT (N = 134 cells, 55 (20 59) cells/mouse [Med (Min Max)]) for event frequency (PVWT = 0.70 (1.21) vs PVAPP = 1.01 (1.62) events/min, X^2^(1) = 0.08, p = 0.78 **(F)** or amplitude (PVWT = 0.68 (0.19) vs PVAPP = 0.72 (0.19) ΔF/F, X^2^(1) = 0.02, p = 0.90) **(G). H** A statistically significant increase in the zero-lag cross-correlation of Ca^2+^ activity between simultaneously recorded PV+ cells was found in PVAPP animals compared with PVWT (PVWT = 0.37 (0.13) vs PVAPP = 0.45 (0.14), X^2^(1) = 5.10, p = 0.024). Violin plots: marker, median; box, IQR; whiskers, lower-upper adjacent values. Box plots show median, IQR and ranges. Descriptive statistics display median and IQR unless otherwise stated. *p < 0.05.

Perisomatic-targeting PV+ interneurons play a key role in generating SWRs^68–70^, and their function is disrupted in several over-expression models of amyloidopathy^13–15^. We thus hypothesised that decreased ripple power in APP mice may be associated with altered PV+ cell function. However, immunohistochemical analysis revealed comparable PV+ cell immunoreactivity between APP and WT mice, suggesting that Aβ pathology did not cause PV+ cell loss (***Supplementary Figure 4E, F***). To measure PV+ cell activity, we generated APP and WT mice expressing Cre recombinase in PV+ neurons (PVAPP and PVWT mice, respectively) and imaged (***Figure 3D***) Cre-dependent GCaMP6f in CA1 PV+ cells under consistent, light anaesthesia (***Supplementary Figure 5***). Most cells (PVAPP = 95.5%; PVWT = 91.9%) exhibited spontaneous somatic Ca^2+^ events indicative of increased neuronal firing^53,71^, that often appeared coordinated between cells in the same field of view (***Figure 3E***). There was no difference in event frequency or amplitude between PVAPP and PVWT interneurons (***Figure 3F, G***). However, Ca^2+^ activity between simultaneously recorded PV+ cells was more correlated for PVAPP cells compared to PVWT (***Figure 3H***). This suggests that Aβ pathology increased coordination of firing rate dynamics between PV+ interneurons without altering activity levels. Since dysfunction in CA1 *oriens-lacunosum moleculare* (O-LM) interneurons has been reported in amyloid over-expression models^72^, we also measured spontaneous Ca^2+^ events from putative O-LM neurons in APP and WT mice expressing Cre recombinase in SST+ interneurons (SSTAPP and SSTWT mice, respectively). However, we observed no change in the frequency or amplitude of spontaneous Ca^2+^ events, or in the correlation of activity between SSTAPP and SSTWT cells (***Supplementary Figure 6***). Thus, hypersynchronous firing of CA1 PV+ cells but not SST+ neurons (which include O-LM cells) may contribute to the observed reduction in ripple power.

### Increased spindle amplitude in the mPFC but no change to the SWO in APP mice

Given that SWRs were found to be impaired during NREM sleep, it was possible that the other neuronal oscillations responsible for long-term memory consolidation, the SWO and sleep spindles, were also affected. The SWO is the temporal pacemaker for systems consolidation and is composed of two distinct states, Up states and Down states (UDS), that reflect periods of synchronous depolarisation and firing followed by subsequent hyperpolarised epochs of relative quiescence^73^. Impairments to the SWO have been observed in both humans with AD^61^ and in mouse models of amyloidopathy^24,25^. However, when we analysed the UDS of the SWO in APP mice, no changes were found relative to WT, nor were there any identifiable deficits to the coupling of the SWO with the gamma oscillations found nested within the Up states (***Supplementary Figure 7***).

The incidence of spindle (11-15 Hz) (***Figure 4A***) oscillations during SWS linearly correlates with subsequent memory performance in humans^41^, with a decrease in the number of detected spindles found in humans with AD^23^. We found a statistically significant increase in the amplitude and corresponding power of spindle events in the mPFC of APP animals relative to WT (***Figure 4B, Cii***), with no change to their duration, frequency, or occurrence (***Figure 4Ci, iii, iv***). PV+ cells in the thalamic reticular nucleus (TRN) are involved in the generation of spindles^74^, while in the mPFC these interneurons are important in the feed-forward inhibition of incoming thalamocortical inputs^75^. Therefore, the density of these interneurons in the TRN and mPFC was assessed using immunohistochemistry. No change in the PV+ cell density was found in the TRN of APP animals (***Supplementary Figure 4A, B***). However, a significant decrease in density was detected in the ACC sub-region of the mPFC (***Supplementary Figure 4C, D***). This decrease in immunoreactivity could potentially reflect decreased capability of PV+ cells to gate incoming TC inputs, producing the increase in spindle amplitude. Thus, it appears that SWR and spindles are altered in APP mice, potentially related to regional changes in the density and activity of PV+ interneurons, whilst the SWO and nested gamma oscillations are spared.

**Figure 4.**
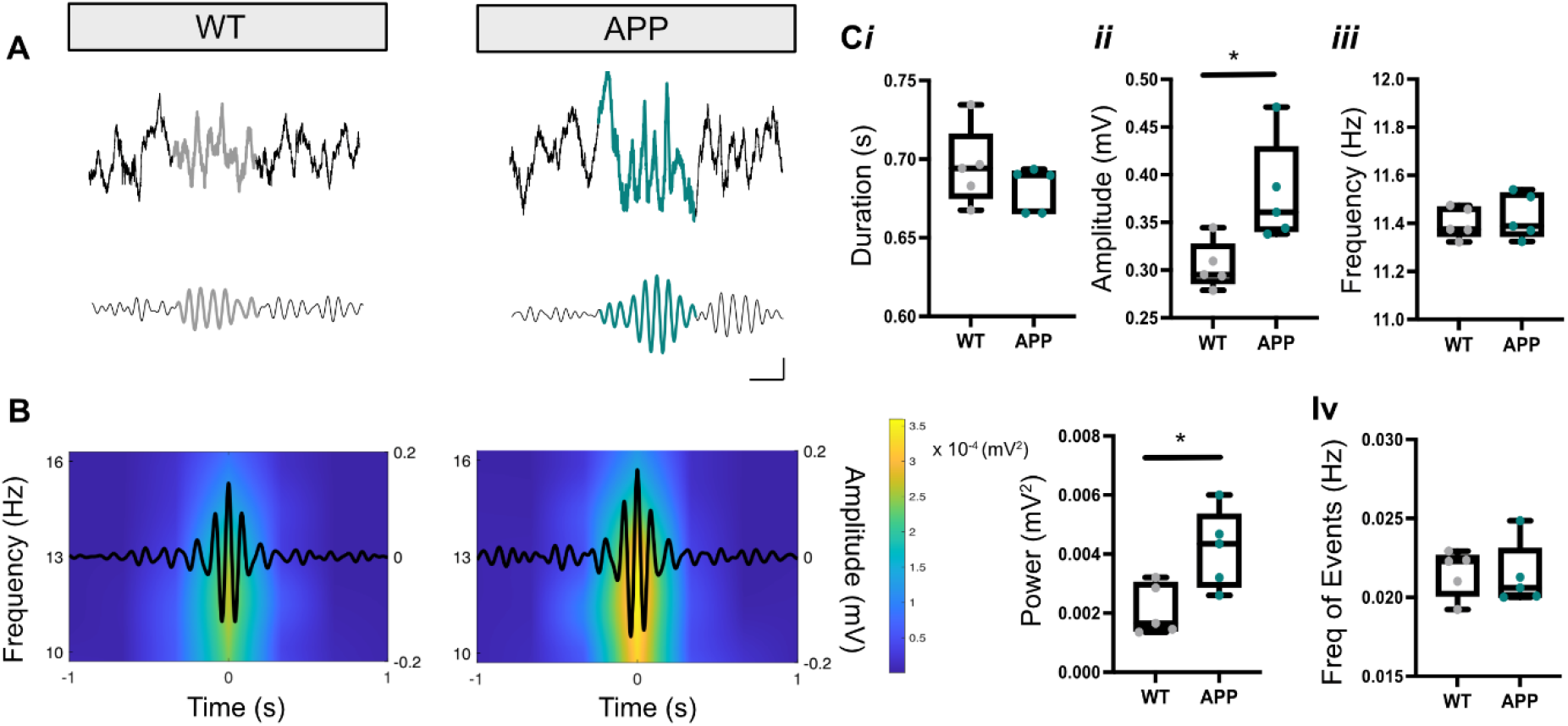
The amplitude and power of mPFC spindles is increased in APP mice. **A** Example raw (top) and 11 – 15 Hz bandpass filtered (bottom) traces of detected spindle events (coloured segments) from WT and APP mice. Scale bars: 250 ms, 200 μV. **B** Example average spindle traces (black lines) overlaid on the corresponding LFP power spectrum showing average spindle power. A statistically significant increase in the average spindle power was found in APP mice compared with WT controls (WT: 0.0017 (0.0014 – 0.0030) vs APP: 0.0043 (0.0029 – 0.0053) mV^2^, U = 2.5, p = 0.03, Mann-Whitney U test). **C** No statistically significant difference was found in APP animals relative to WT for the average spindle duration (WT: 0.69 (0.68 – 0.0.72) vs APP: 0.69 (0.67 – 0.69) s, U = 6, p = 0.22, Mann-Whitney U test) ***(i)*** or frequency (WT: 11.38 (11.35 – 11.47) vs APP: 11.39 (11.35 – 11.53) Hz, U = 10, p = 0.69, Mann-Whitney U test) ***(iii)*** but a statistically significant increase in the average spindle amplitude was found in APP mice compared with WT controls (WT: 0.30 (0.29 – 0.33) vs APP: 0.36 (0.34 – 0.43) mV, U = 2, p = 0.03, Mann-Whitney U test) ***(ii)***. No statistically significant difference was found in APP animals relative to WT for the incidence of detected spindle events (WT: 0.022 (0.020-0.023) vs APP: 0.021 (0.020 – 0.023) Hz, U = 10, p = 0.69, Mann-Whitney U test) ***(iv)***. Box plots show median, IQR and ranges. Descriptive statistics display median and IQR. *p < 0.05.

### Temporal communication between NREM sleep-associated neuronal oscillations is intact in APP mice

While the SWO, spindles and SWR play important functional roles in memory processing within the hippocampal-mPFC circuit, it is their temporal coupling that is thought to drive systems consolidation^20,21^. Cortical spindles occur at the beginning of SWO Up states^46,76^ while SWR occur towards the end^77^. This coupling could be clearly seen in our recordings when plotting the power of mPFC spindles and hippocampal SWR relative to the SWO trace (***Figure 5A, D***). To assess the coupling strength between these oscillations, several different parameters were measured. No significant change was found in the phase-amplitude coupling (PAC) of the SWO with cortical spindles when comparing APP with WT mice (***Figure 5B***). Additionally, there was no significant change in the time lag between the peak of the SWO Down states and the peak of cortical spindle events, nor was there a change in the percentage of SWO events that were coupled to a cortical spindle (***Figure 5C***). Furthermore, no statistically significant change was found in APP animals relative to controls when looking at the PAC the SWO with ripples (***Figure 5E***) and when comparing the time lag between the peak of the Down states with the peak of the ripple events (***Figure 5F***). However, a trending increase in the percentage of SWO events coupled to ripples was observed in APP animals compared with WT controls (***Figure 5F***). This is consistent with the trend towards increased ripple incidence reported in ***Figure 3Civ***.

**Figure 5.**
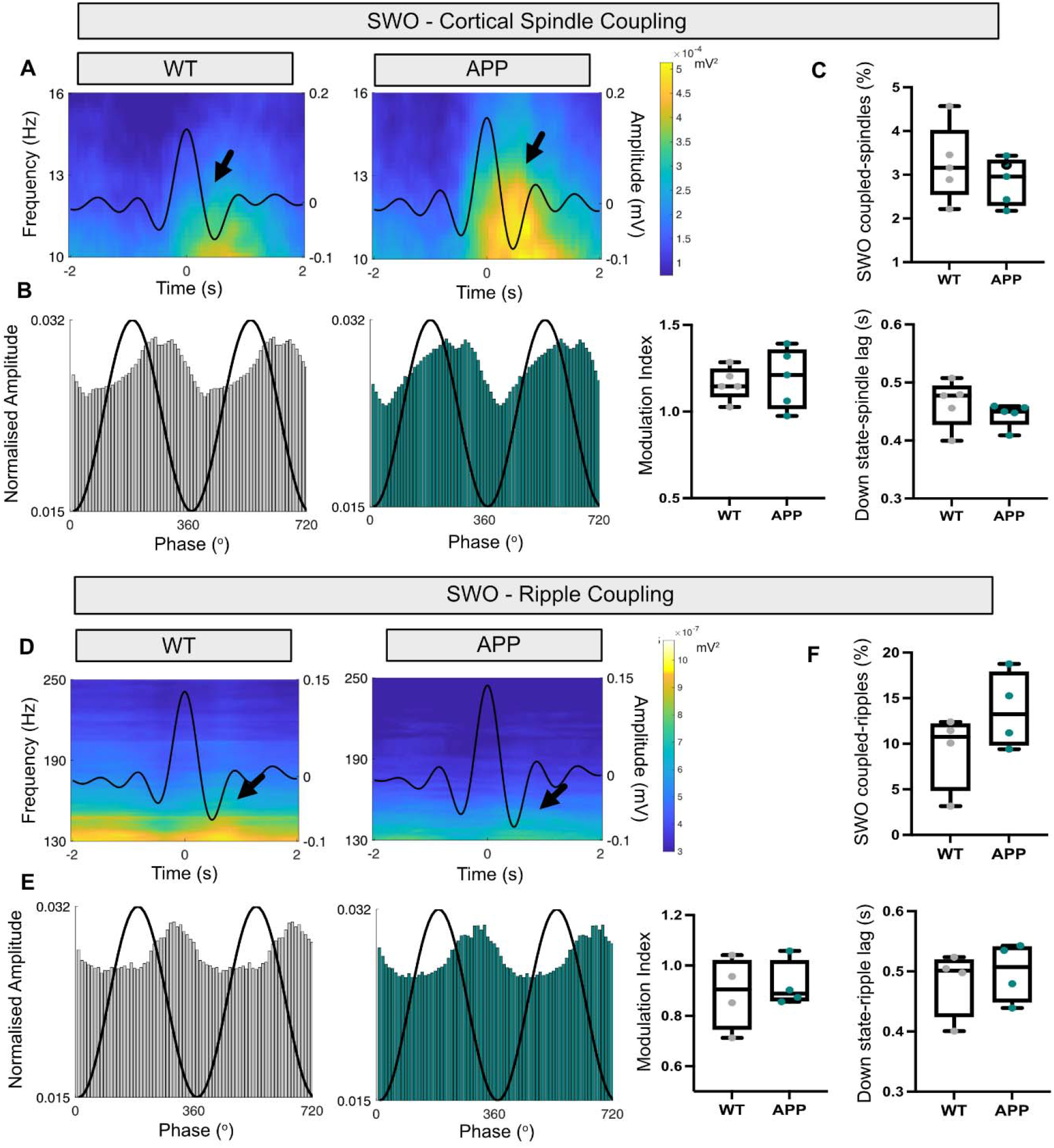
Coupling of UDS with mPFC spindles and hippocampal SWR is not altered in APP mice. **A** Example average UDS traces (black lines) overlaid on the corresponding LFP power spectrum showing the power of UDS-locked mPFC spindle oscillations for both WT and APP mice. Spindles can be seen locked to SWO Up states (black arrows). **B** No statistically significant difference in the PAC of UDS with mPFC spindle oscillations was found in APP animals relative to WT (WT: 1.15 (1.09 – 1.24) vs APP: 1.21 (1.02 – 1.36) MI, U = 10, p = 0.65, Mann-Whitney U test). Shown are example histograms of the change in normalised spindle amplitude (bars) over the phase of the UDS (black line) for both WT (left) and APP (right) mice. **C** No statistically significant difference was found in the time lag between the peak of the Down state to the peak of the mPFC spindle event (WT: 0.48 (0.43 – 0.49) vs APP: 0.45 (0.43 – 0.46) s, U = 7, p = 0.31, Mann-Whitney U test) or in the percentage of UDS coupled to a mPFC spindle event (WT: 3.16 (2.55 – 4.01) vs APP: 2.96 (2.30 – 3.33) %, U = 9, p = 0.54, Mann-Whitney U test). **D** Example average UDS traces (black lines) overlaid on the corresponding LFP power spectrum showing the power of UDS-locked ripple oscillations for both WT and APP mice. Ripples can be seen locked to SWO Up states (black arrows). **E** No statistically significant difference in the PAC of UDS with ripple oscillations was found between in APP animals compared with WT controls (WT: 0.90 (0.75 – 1.02) vs APP: 0.89 (0.86 – 1.02) MI, U = 6, p = 0.68, Mann-Whitney U test). Shown are example histograms of the changes to the normalised ripple amplitude (bars) over the phase of the UDS (black line) for both WT (left) and APP (right) mice. **F** No statistically significant difference was found in the time lag between the peak of the Down state to the peak of the ripple event (WT: 0.50 (0.42 – 0.52) vs APP: 0.51 (0.44 – 0.54) s, U = 6, p = 0.68, Mann-Whitney U test) but trending increase was found in APP mice compared with WT controls when looking at the percentage of UDS coupled to ripples (WT: 10.76 (4.89 – 12.15) vs APP: 13.23 (9.86 – 17.88) %, U = 5, p = 0.48, Mann-Whitney U test). Graphs show median and ranges. Box plots show median, IQR and ranges. Descriptive statistics display median and IQR.

Next, we investigated the coupling of spindles with SWRs. Cortical spindles are initiated before hippocampal ripples^78–80^ (***Figure 6A***); it is thought that spindles prime cortical neurons for plasticity-related changes by increasing dendritic Ca^2+ 47^ just prior to information transfer from the hippocampus during a ripple^65,66^. A breakdown in this coupling could therefore have disastrous consequences for memory consolidation. Nonetheless, no statistically significant change was found in APP animals relative to WT controls in the time lag between the start of cortical spindles and hippocampal ripples (***Figure 6B***), nor in the percentage of cortical spindles coupled to ripples (***Figure 6C***). Finally, no significant difference was found in the occurrence of SWO events with both spindles and ripples when comparing APP animals with WT (***Figure 6D,E***). Therefore, despite changes occurring to local spindle and SWR events in APP mice, the ability of the mPFC-hippocampal circuit to control the temporal precision of neural activity remains intact.

**Figure 6.**
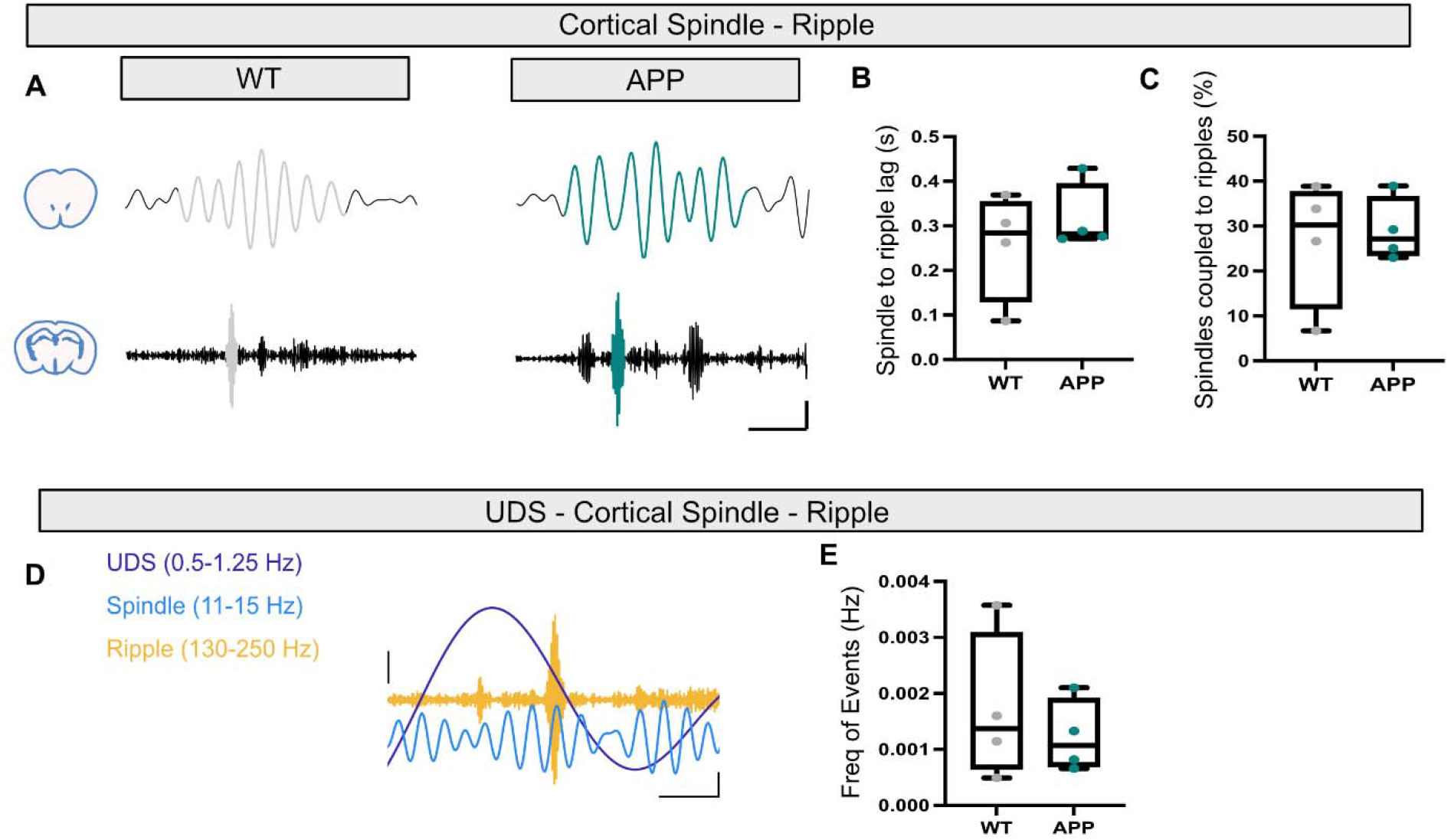
UDS-spindle-ripple coupling is intact in APP mice. **A** Examples of 11-15 Hz bandpass filtered traces of spindle events in the mPFC (top) coupled to 130 – 250 Hz bandpass filtered CA1 ripple events (bottom) from WT and APP mice (coloured segments). Scale bars: 200 ms, 100 μV. **B** No statistically significant difference was found in the time lag between the start of mPFC spindle events to the start of CA1 ripple events (WT: 0.28 (0.13 – 0.35) vs APP: 0.28 (0.27 – 0.39) s, U = 6, p = 0.68, Mann-Whitney U test). **C** No statistically significant difference was found in the percentage of mPFC spindles coupled to CA1 ripples (WT: 30.24 (11.69 – 37.61) vs APP: 27.16 (23.51 – 36.52) %, U = 8, p = 0.99, Mann-Whitney U test). **D** Example of bandpass filtered traces showing the three cardinal oscillations occurring together in a WT mouse. UDS (0.5 – 1.25 Hz, purple), spindles (11 – 15 Hz, blue) and ripples (130 – 250 Hz, orange). Scale bars: 200 ms, 30 μV (right – UDS and spindle), 100 μV (left - ripple). **E** No statistically significant difference in the frequency of the three cardinal oscillations occurring together (WT: 0.00066 (0.00065 – 0.00308) vs APP: 0.0011 (0.00070 – 0.00190) %, U = 7, p = 0.88, Mann-Whitney U test). Box plots show median, IQR and ranges. Descriptive statistics display median and IQR.

## Discussion

This report is the first to assess rhythmic network activity across different brain states in the App^NLG-F/NL-G-F^ mouse model. It is also the first study to comprehensively analyse hippocampal-mPFC circuit function during NREM sleep in a mouse model of amyloidopathy, although others have made enquiries along similar lines^81,82^. Across awake, REM and NREM brain states the power of gamma oscillations was unperturbed in both the mPFC and CA1 region of the hippocampus. However, during NREM sleep changes to cortical spindles as well as hippocampal ripples were observed in App^NL-G-F/NL-G-F^ mice at sixteen months of age. These changes to NREM sleep-associated neuronal oscillations measured in local networks were not accompanied by deficits to the temporal coupling between different oscillations throughout the hippocampal-mPFC circuit. Finally, PV+ interneurons in CA1 were found to exhibit a higher degree of synchronous activity, and a decrease in PV+ interneuron density was identified in the ACC. Taken together, these results indicate that, prior to the onset of spatial and contextual memory impairments, Aβ pathology changes PV+ interneuron-mediated inhibition in local neural circuits, likely disrupting local oscillatory dynamics specific to NREM sleep without disrupting long-range communication between hippocampus and mPFC. These deficits may be a harbinger of subsequent breakdown of hippocampal and mPFC function in AD.

### Gamma oscillations across brain states are unimpaired

During active exploration, gamma oscillations show reduced power within the cortex and hippocampus of first-generation mouse models of amyloidopathy (i.e. mice overexpressing human APP with familial AD mutations)^13,14,63^. Therefore, it was hypothesised that similar disruptions to gamma oscillations would be found during active exploration in App^NL-G-F/NL-G-F^ mice, as well as during REM and NREM sleep. However, no differences were identified in the power of either low or high frequency gamma oscillations in either wake, REM or NREM sleep, across both the mPFC and CA1 region of the hippocampus. This is in contrast to a previous study that discovered evidence for impaired CA1 high frequency gamma oscillations during spatial navigation in the App^NL-G-F/NL-G-F^ model between seven-thirteen months of age^62^. Possible explanations for this discrepancy could be our preferred use of wildtype littermate controls compared to non-littermate control mice used in the prior study^62^. Despite no statistically significant differences, a trending decrease in low and high gamma oscillatory power was observed in CA1 across brain states, potentially signalling the beginning of gamma oscillation breakdown in this brain region at sixteen-months. This is in comparison to first-generation amyloidopathy models that can display deficits in gamma oscillations as early as 4 months of age^13^. It is therefore possible that when mutated App is expressed at more physiological levels, a more prolonged exposure to amyloid pathology is needed for gamma oscillation impairments to manifest.

### An increase in mPFC spindle amplitude and a decrease of PV+ cell density in the ACC

An increase in the amplitude and power of cortical spindles was identified in APP animals compared to WT controls. Given that PV+ cells within the anterior TRN help generate spindles^37^ and that cortical PV+ interneuron function is documented to be impaired in first-generation mouse models of AD^13,14^, the density of these neurons was quantified. No difference was found between genotypes in the TRN, although this does not necessarily rule out impaired PV+ cell function. Additionally, there was a statistically significant decrease in the density of PV+ cells in the ACC. PV+ cells in the mPFC help gate incoming excitatory TC input during spindles, and prevent action potential propagation, localising increased Ca^2+^ to the dendrites to facilitate plasticity-related changes^75^. A potential loss of these interneurons could impair this function and contribute to the increase in spindle amplitude. If this is the case, the failure of PV+ cells to regulate somatic and dendritic excitability could have disastrous ramifications for plasticity. However, these hypotheses are speculative and need to be investigated.

### Disrupted hippocampal ripples and increased coordination of PV+ cell activity

Ripple events recorded in CA1 *Str.P* exhibited a statistically significant reduction in power but, interestingly, no change in amplitude. Power was calculated from the short-time FFT of the whole event whereas amplitude was the difference between the highest peak and lowest trough, typically located in the centre of the ripple event. This suggests disrupted spike-time dynamics occurring elsewhere in the ripple oscillation. Indeed, CA1 PCs display impaired spiking during fast gamma oscillations in the App^NL-G-F/NL-G-F^ mouse model^62^. Both fast gamma oscillations and ripples require the recruitment of PV+ interneurons to modulate PC firing^70,83^, and PV+ interneuron spiking is reported to be impaired in first-generation mouse models of amyloidopathy^13,14^.

Ca^2+^ imaging revealed increased coordination of PV+ interneuron activity in CA1 in App^NL-G-F/NL-G-F^ mice, yet these cells exhibited no change in their activity levels or density. This implies that compensatory alterations may occur to the CA1 network, potentially as a result of altered neuronal activity^6,62,84^. Moreover, the increased coordination of PV+ cell activity could explain the observed decrease in ripple power. PV+ cells are involved in the pacing the ripple oscillation^70^ and follow the spiking of PCs in the ripple trough^69^. An increase in firing coordination could decrease the excitation-inhibition balance during each ripple cycle, resulting in a reduction in power and potential changes to PV+ interneuron – PC firing dynamics. This not only has the potential to disturb processes such as spike-time-dependant plasticity, but it could affect the recruitment of neurons for the reactivation of previously encoded information^85^, potentially interfering with consolidation. Future experiments recording the activity of both PCs and PV+ cells during SWRs in App^NL-G-F/NL-G-F^ mice will give a clearer indication of how changes in PV+ interneuron activity contribute to dysfunctional oscillatory dynamics. Moreover, given the reduced power of ripples, it would be interesting for future experiments to look at the spiking activity of neurons in the mPFC during a ripple event, to determine if the information contained within the ripple is being effectively transmitted to the cortex^65,66^.

### The SWO and temporal communication within the hippocampal – mPFC circuit is spared

Although impairments were seen in both cortical spindles and hippocampal ripples, the temporal coupling between cardinal oscillations remained intact in sixteen-month-old APP mice. SWO coupling to cortical spindles was unaffected, suggesting that the TC feedback loop that controls their temporal communication is spared^45^. Further evidence of spared communication between brain regions is our observation of intact coupling between the SWO and hippocampal ripples, cortical spindles with ripples, as well as the occurrence of all three cardinal oscillations together. This lack of coupling deficits is in line with no observable impairments to the SWO, which is considered the temporal pacemaker of the hippocampal – mPFC circuit during NREM sleep. Dysfunction of the SWO is reported in several over-expression mouse models of amyloidopathy^24–26^; therefore, like gamma oscillations, it is possible that amyloidopathy alone has no effect upon the SWO when mutated App is expressed at more physiological levels.

Taken together, these results suggest hippocampal-mPFC long-range communication is spared, with the changes occurring to the local oscillations constituting the first features of circuit dysfunction in App^NL-G-F/NL-G-F^ mice. However, it is important to note that learning induces changes to oscillatory activity in the systems consolidation circuit: increased slow wave activity, spindle and ripple density, as well as an increase in spindle-ripple coupling are reported^41^. In some studies, Aβ-induced deficits to SWR dynamics were only identified during post-learning sleep following a hippocampus-dependant memory task, as opposed to during baseline conditions^86,87^. It is important to assess hippocampal-mPFC circuit function in baseline conditions without prior cognitive loading, however, as learning-induced changes could mask subtle neurophysiological deficits as biomarkers of AD. Yet, it is possible Aβ-induced deficits are more pronounced during active consolidation of hippocampus-dependant memories. Therefore, repeating experiments under these conditions may produce more striking deficits.

### Relevance to the human condition

Gamma oscillations are considered a biomarker of network dysfunction in AD. Yet, studies conducted in humans produce varying results. Both increased and decrease gamma oscillation power have been identified, with no clear relationship between power, brain state or pathological state of the patients^61,88^. These wide-ranging effects in humans coupled with the lack of statistically significant differences in gamma oscillation power in the present study suggest that impairments to these oscillations may not reflect an early or reliable instance of network dysfunction in AD. In fact, according to the data presented in this study, alterations to cortical spindle and hippocampal SWRs precede gamma oscillation breakdown and may potentially be a better marker of early network dysfunction as a result of amyloid pathology in AD.

The development of amyloid pathology is thought to precede and contribute to the cascade of pathophysiological impairments seen in AD^89^. Additionally, sleep disruptions are common in the early stages of the disease and are thought to be a catalyst for further pathophysiological impairments^55^. It is therefore possible that the App^NL-G-F/NL-G-F^ mouse model of amyloidopathy reflects these early stages of AD progression, making it an ideal model for identifying biomarkers of early network dysfunction and investigating the initial causes of circuit breakdown. It is important that research into these early preclinical stages of AD is conducted, as it is likely that treatments targeting these stages are more effective^90^.

