## Supplementary material for "Alterations to parvalbumin-expressing interneuron function and associated network oscillations in the hippocampal – medial prefrontal cortex circuit during natural sleep in App^NL-G-F^ mice"

### Supplementary Figures

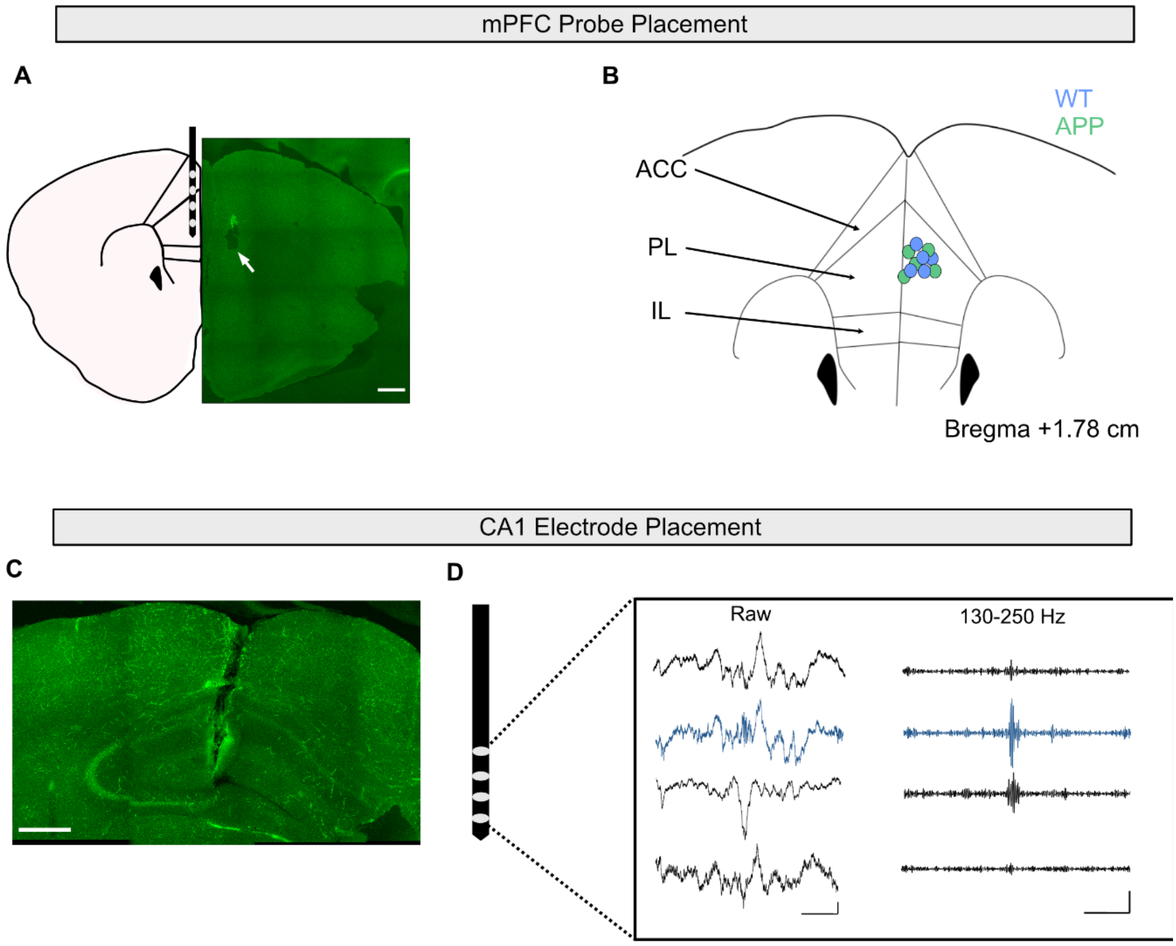


**Supplementary Figure 1 | Verification of electrode placement. A** Schematic of mPFC sub-regions with silicon probe placement (left) and an example image of an electrolytic lesion at the bottom electrode site (right). White arrow points to the lesion. Scale bar: 500 μm. **B** Schematic summarising the placement of mPFC silicon probes in WT (blue) and APP (green) animals as verified by histological markings. Dots represent tip of silicon probe. **C** Image showing a tract left by a silicon probe in the CA1 region of the hippocampus. Scale bar: 500 μm. **D** Raw and 130-250 Hz bandpass filtered traces displaying a SWR event recorded using a 4-channel silicon probe electrode. In blue is a trace recorded in the *Str.P* layer of CA in a WT mouse. SWRs were used to identify the locations of electrodes across layers of CA1. Scale bars: 200 ms, 0.1 mV. Representative images are from WT animals were immunohistochemically stained with anti-$A\beta_{1-42}$.

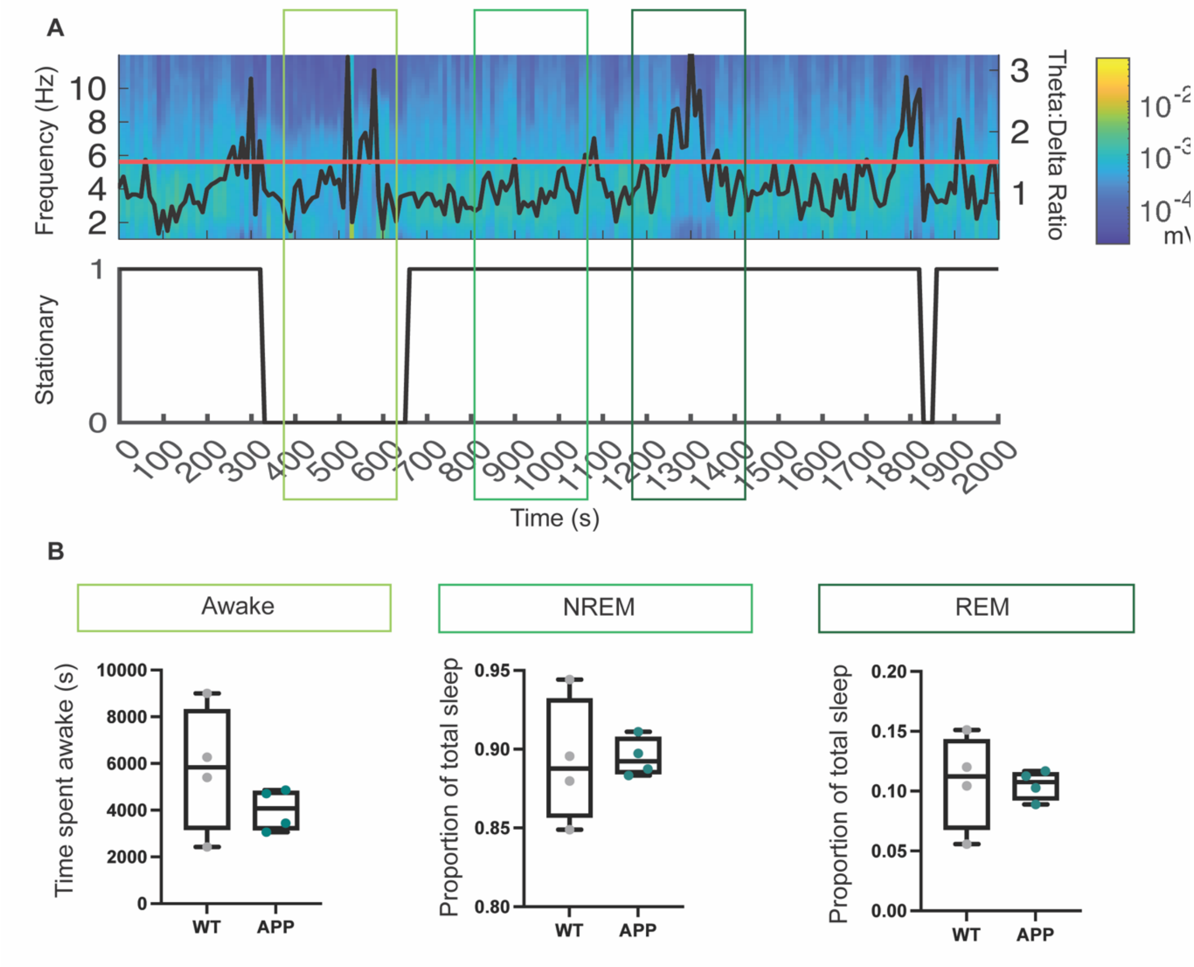


**Supplementary Figure 2 | WT and APP mice spend comparable durations in awake, REM and NREM states. A** (Above) Spectrogram showing power changes over time within the combined UDS and delta band (1-4 Hz) and theta (6-12 Hz) band. Black line represents the theta:delta power ratio over time. Horizontal red line shows the threshold for distinguishing periods of awake/REM sleep from NREM sleep (median + 1SD). (Below) Corresponding manually-scored movement of the animal over time, taken as a binary measure, with 1 showing no movement and 0 showing movement. Portions of the time series are highlighted with coloured boxes to signify periods of detected awake states, NREM and REM sleep. B WT and APP mice spent comparable amount of time in awake states (WT: 5835 (3137-8318) vs APP: 4080 (3158-4823) s, U = 4 $p = 0.34$, Mann-Whitney U test), NREM sleep (WT: 0.89 (0.86-0.93) vs APP: 0.89 (0.88-0.91) proportion of total sleep, U = 6, $p = 0.68$, Mann-Whitney U test) and REM sleep (WT: 0.11 (0.07-0.14) vs APP: 0.1 (0.09-0.12) proportion of total sleep, U = 6, $p = 0.68$, Mann-Whitney U test). Box plots show median, IQR and ranges. Descriptive statistics display median and IQR.

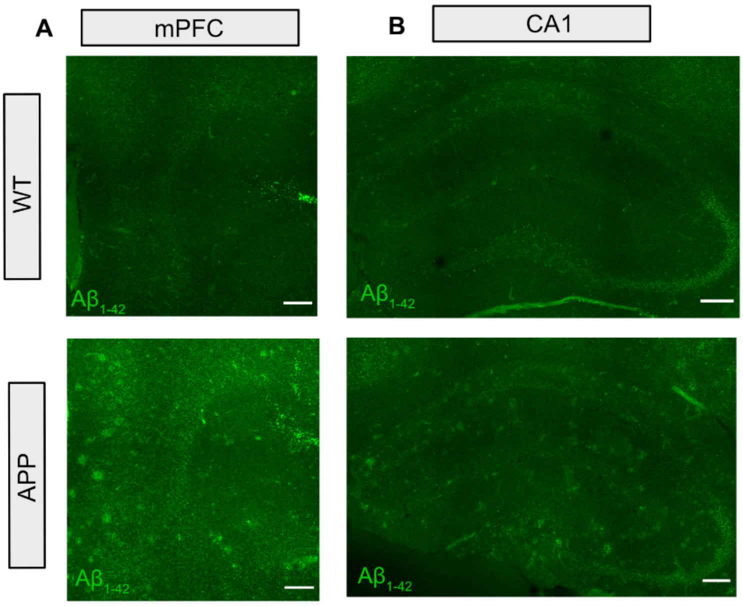


**Supplementary Figure 3 | Extensive A$\beta_{1\text{-}42}$ pathology found in the mPFC and hippocampus of sixteen months APP mice.** Representative images of A$\beta_{1\text{-}42}$ plaques immunolabelled within the medial prefrontal cortex (mPFC) **(A)** and hippocampus **(B)** of APP mice. Note the absence of plaque labelling in littermate WT mice. Scale bar: 200 μm.

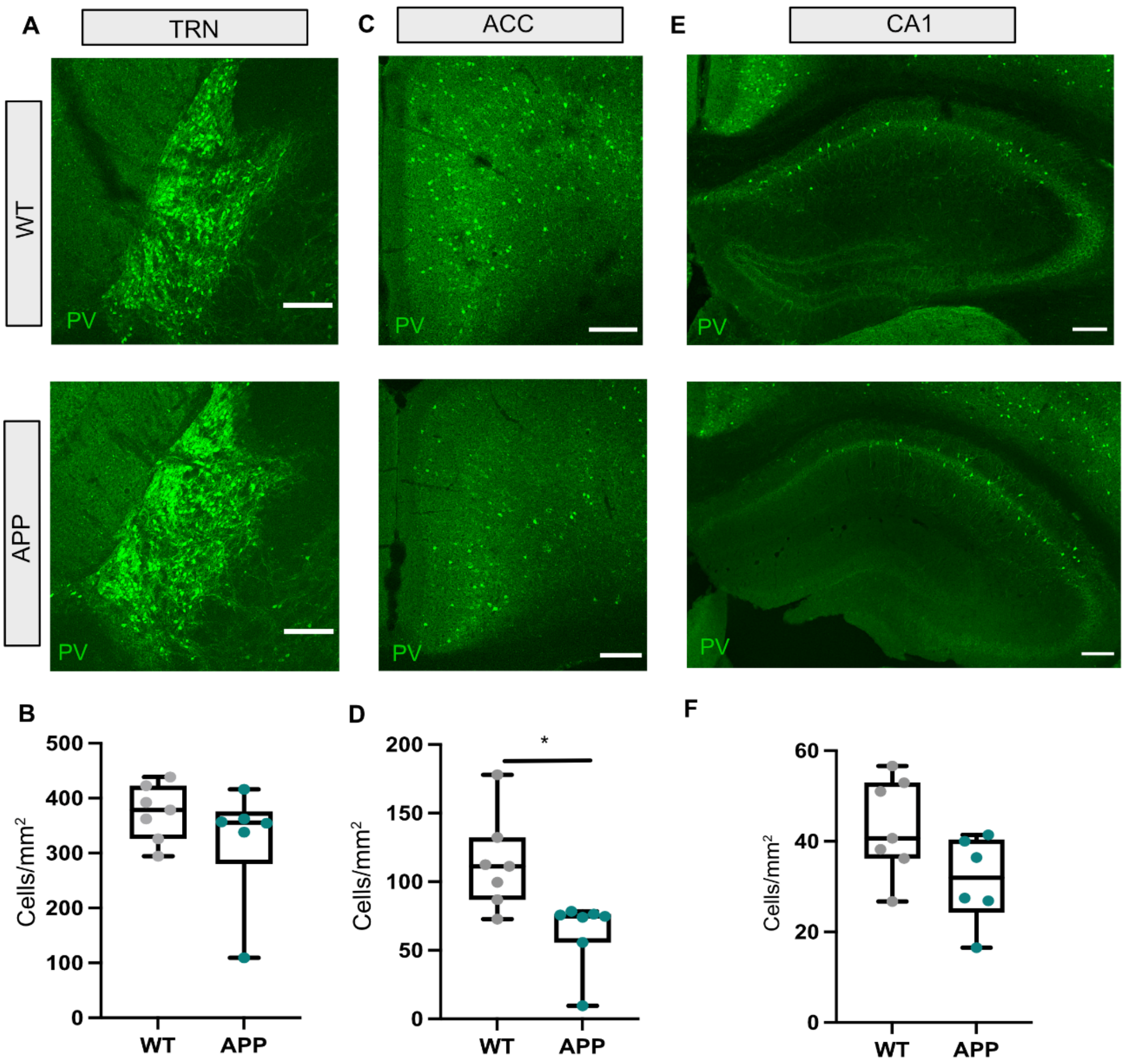


**Supplementary Figure 4 | Decreased PV-expressing (PV+) cell density in the ACC sub-region of the mPFC.** Representative images of PV+ interneurons within the thalamic reticular nucleus (TRN) **(A)**, anterior cingulate cortex (ACC) **(C)** and hippocampus **(D)** of WT and APP mice. Scale bar: 200 μm. **B** No statistically significant difference in PV+ cell immunoreactivity in the TRN between genotypes (WT: 378.7 (326.1-422.5) vs APP: 355.5 (280.8-375.7) Cell/mm$^2$, U = 13.5, $p$ = 0.31, Mann-Whitney U test). **D** A statistically significant decrease in PV+ cell immunoreactivity in the ACC of APP animals (WT: 111.30 (86.96-132.20) vs APP: 74.82 (55.71-76.35) cells/mm$^2$, U = 5, $p$ = 0.01, Mann-Whitney U test). **F** No statistically significant difference in PV+ cell immunoreactivity in CA1 between genotypes WT: 40.70 (36.22-52.94) vs APP: 31.95 (24.30-40.36) cells/mm$^2$, U = 11, $p$ = 0.18, Mann-Whitney U test). Box plots show median, IQR and ranges. Descriptive statistics display median and IQR.

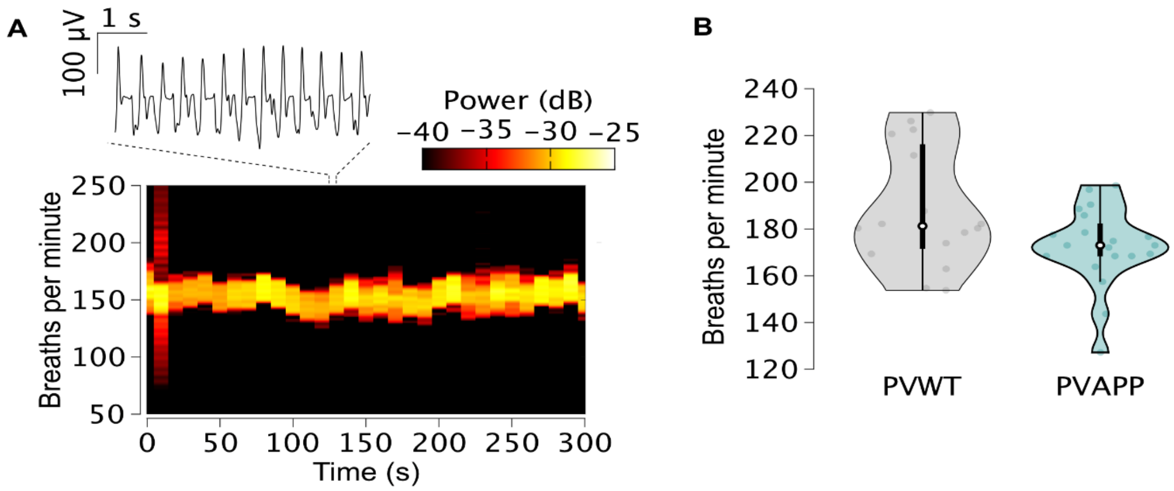


**Supplementary Figure 5 | Consistent recording conditions for throughout in vivo $Ca^{2+}$ imaging experiments. A** Example spectrogram plotting a mouse's breathing rate throughout a 300 s 2-photon imaging experiment. The inset trace (above) illustrates a 5 s record from the piezo breathing sensor, where each spike corresponds to an individual breath (scale bar: 100 µV, 1 s). Note that a stable rate around 150 breaths per minute was maintained throughout the experiment. **B** A comparable breathing rate was maintained across experiments (PVWT: 181.3 (47.8) vs PVAPP: 173.0 (15.6), $F(1,34) = 1.86$, $P = 0.18$). Descriptive statistics display median and IQR.

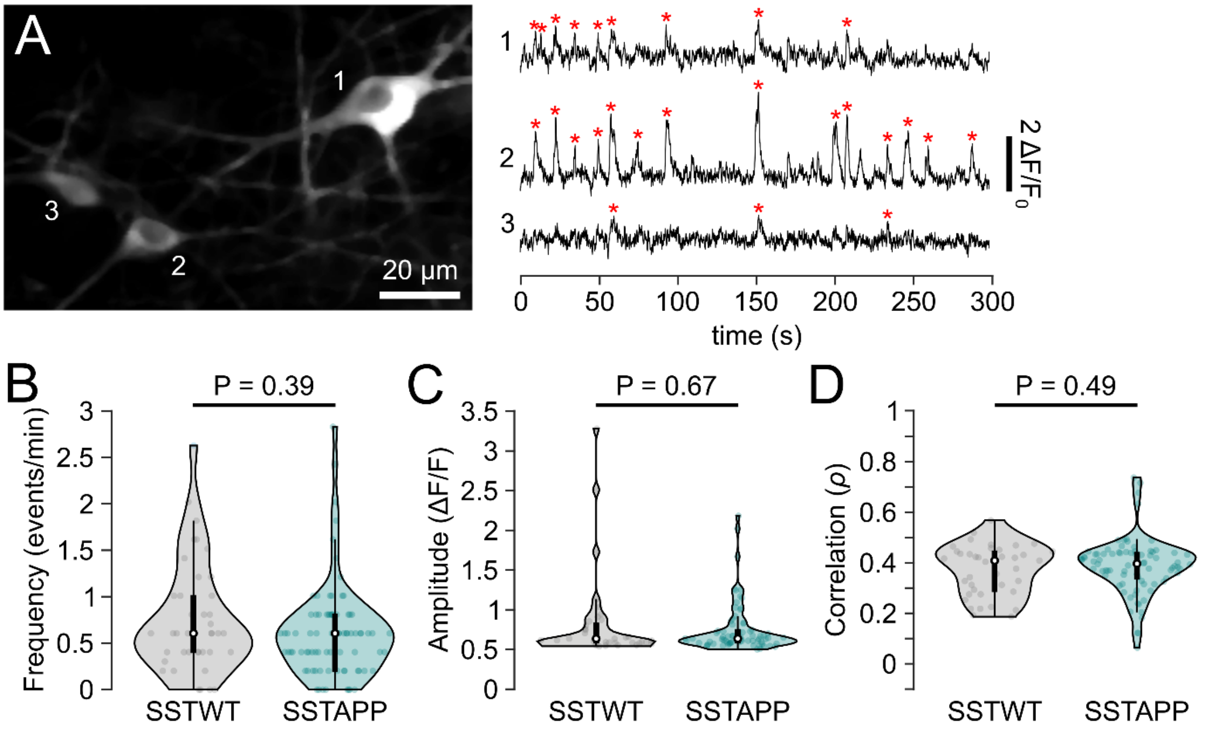


**Supplementary Figure 6 | Amyloidopathy does not disrupt the activity of putative O-LM interneurons in APP mice. (A)** Left: Two-photon image of three representative GCaMP6s-expressing cells in an SSTAPP mouse. Right: GCaMP6f fluorescence traces (ΔF/F) recorded from each cell. Red stars denote detected spontaneous $Ca^{2+}$ events. **B-D** No statistically significant change was identified for SSTAPP cells (N = 82 cells, 22.5 (13 24) cells/mouse [Med (Min Max)]) compared to SSTWT (N = 39 cells, (10 29) cells/mouse (Min Max) for event frequency (SSTWT = 0.61 (0.61) vs SSTAPP = 0.61 (0.56) events/min, $X^2(1,121) = 0.76$, p = 0.39) **(B)**, event amplitude (SSTWT = 0.64 (0.24) vs SSTAPP = 0.64 (0.16) ΔF/F, $X^2(1,121) = 0.18$, p = 0.67) **(C)**, or the zero-lag cross-correlation of $Ca^{2+}$ activity between simultaneously recorded cells (SSTWT = 0.41 (0.15) vs SSTAPP = 0.40 (0.10), $X^2(1,121) = 0.47$, p = 0.49) **(D)**. Violin plots: marker, median; box, IQR; whiskers, lower-upper adjacent values. Descriptive statistics report the median and IQR unless otherwise stated.

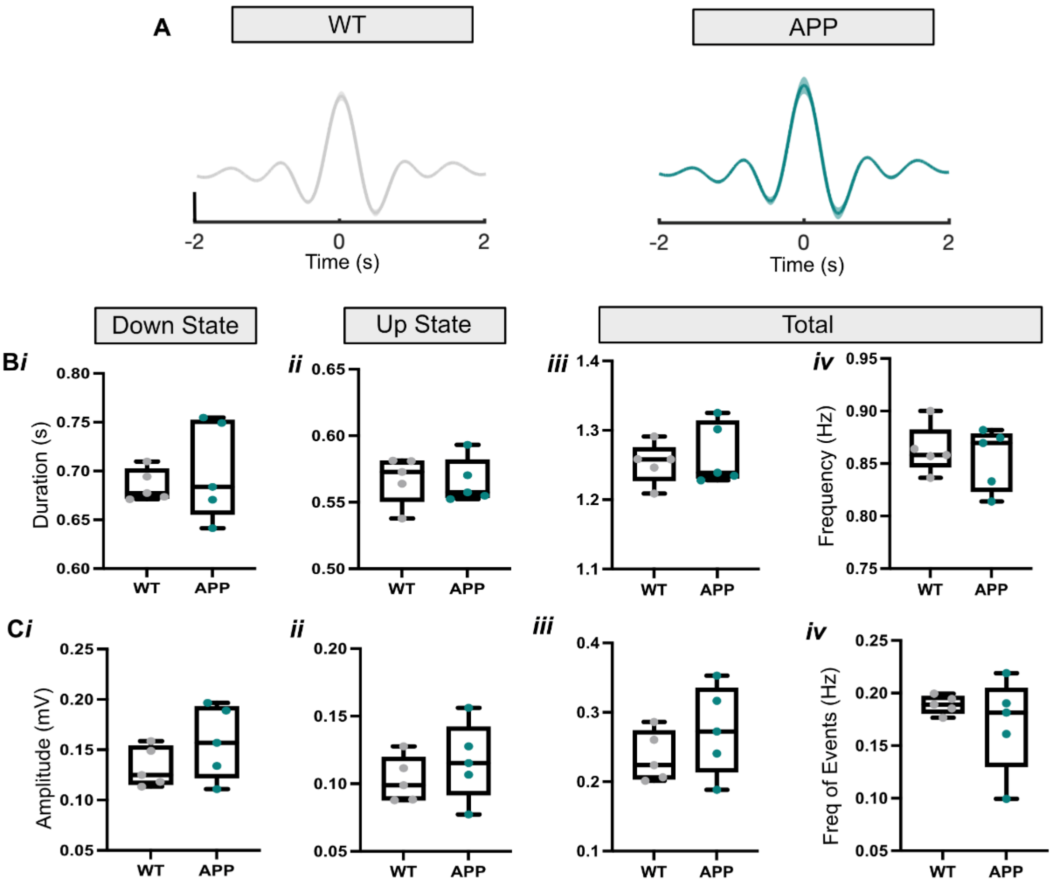


**Supplementary Figure 7 | UDS dynamics are not altered in APP animals compared with WT controls. A** Average ± SEM traces of detected UDS, centred around the peak of the Down state, for both WT and APP mice. Scale bar: 50 µV. **B** No statistically significant difference was found in APP animals compared with WT for the average Down state duration (*WT:* 0.67 (0.67-0.70) *vs APP:* 0.68 (0.66-0.75) s, $U$ = 12, $p$ = 0.99, Mann-Whitney U test) ***(i)***, Up state duration (*WT:* 0.57 (0.55-0.58) *vs APP:* 0.56 (0.55-0.58) s, $U$ = 10, $p$ = 0.69, Mann-Whitney U test) ***(ii)***, total duration (*WT:* 1.26 (1.23-1.28) *vs APP:* 1.24 (1.23-1.31) s, $U$ = 12, $p$ = 0.99, Mann-Whitney U test) ***(iii)***, or UDS frequency (*WT:* 0.86 (0.85-0.88) *vs APP:* 0.87 (0.82-0.88) Hz, $U$ = 12, $p$ = 0.99, Mann-Whitney U test) ***(iv)***. **C** No statistically significant difference was found between genotypes for the average Down state amplitude (*WT:* 0.13 (0.12-0.15) *vs APP:* 0.16 (0.12-0.19) mV, $U$ = 8, $p$ = 0.42, Mann-Whitney U test) ***(i)***, Up state amplitude (*WT:* 0.10 (0.09-0.12) *vs APP:* 0.12 (0.09-0.14) mV, $U$ = 8 $p$ = 0.42, Mann-Whitney U test) ***(ii)***, total amplitude (*WT:* 0.22 (0.20-0.27) *vs APP:* 0.27 (0.21-0.33) mV, $U$ = 8, $p$ = 0.42, Mann-Whitney U test) ***(iii)*** nor number of detected UDS events (*WT:* 0.18 (0.18-0.20) *vs APP:* 0.18 (0.13-0.20) Hz, $U$ = 9, $p$ = 0.54, Mann-Whitney U test) ***(iv)***. Box plots show median, IQR and ranges. Descriptive statistics display median and IQR.
